# Detection of cooperatively bound transcription factor pairs using ChIP-seq peak intensities and expectation maximization

**DOI:** 10.1101/120113

**Authors:** Vishaka Datta, Rahul Siddharthan, Sandeep Krishna

## Abstract

Transcription factors (TFs) often work cooperatively, where the binding of one TF to DNA enhances the binding affinity of a second TF to a nearby location. Such cooperative binding is important for activating gene expression from promoters and enhancers in both prokaryotic and eukaryotic cells. Existing methods to detect cooperative binding of a TF pair rely on analyzing the sequence that is bound. We propose a method that uses, instead, only ChIP-seq peak intensities and an expectation maximization (CPI-EM) algorithm. We validate our method using ChIP-seq data from cells where one of a pair of TFs under consideration has been genetically knocked out. Our algorithm relies on our observation that cooperative TF-TF binding is correlated with weak binding of one of the TFs, which we demonstrate in a variety of cell types, including *E. coli, S. cerevisiae* and *M. musculus* cells. We show that this method performs significantly better than a predictor based only on the ChIP-seq peak distance of the TFs under consideration. This suggests that peak intensities contain information that can help detect the cooperative binding of a TF pair. CPI-EM also outperforms an existing sequence-based algorithm in detecting cooperative binding. The CPI-EM algorithm is available at https://github.com/vishakad/cpi-em.

## 1 Introduction

Transcription factors (TFs) regulate the transcription of a set of genes by binding specific regulatory regions of DNA. The magnitude of the change in transcription caused by a TF depends in part on its affinity to the bound DNA sequence. Some times, it is possible that a second TF binding a nearby sequence increases the first TF’s binding affinity. In this case, the two TFs are said to cooperatively or combinatorially bind DNA [1]. The cooperative binding of transcription factors at enhancers and promoters is known to strongly increase gene expression [2, 3, 4, 5]. The presence of cooperativity has been used to explain the rapid rate of evolution of TF binding sites in multicellular organisms [6].

The role of cooperative binding in protein complex assembly has been extensively studied and computational methods have been proposed to detect such interactions within genomes [7, 8, 9]. In these studies, cooperativity results in the oligomerization of proteins after they bind DNA through protein-protein contacts. In such TF pairs, this typically occurs only when their binding sites are at a particular distance from each other. Earlier theoretical methods have successfully detected many such instances of cooperatively bound TF pairs [10, 11, 12, 13, 14, 15, 16, 1, 17]. The input to these methods is a set of sequences bound by both TFs under investigation. These methods scan these sequences for closely spaced binding sites of both TFs, using position weight matrix (PWM) models of each TF [18], and predict the distance between the binding sites at which cooperative interactions can occur.

However, many TF pairs can cooperatively bind DNA even if the distance between their binding sites is changed [19], and need not form protein-protein contacts upon binding DNA [20, 21]. The strength of the cooperative effect in these cases can depend on the distance between the binding sites [21]. Such a distance-independent cooperative interaction can arise from a mechanism such as assisted binding [22], where a TF, say A, that is already bound to DNA increases the affinity of nearby binding site towards a second TF, say B. Such a cooperative interaction may be asymmetric in nature i.e., a TF A may be able to assist a TF B in binding DNA, but not vice versa [22]. The sequence that links the two binding sites can also modulate the cooperative effect. For instance, nucleotide substitutions in the sequence linking binding sites of the transcription factors Sox2 and Pax6 were found to convert a cooperatively bound enhancer sequence in the *D. melanogaster* genome to a non-cooperatively bound one [23].

An important consequence of these findings is that a pair of TFs that cooperatively bind DNA at one genomic region may not bind cooperatively at a different genomic region due to differences in the binding site arrangement between both regions. For such TF pairs, it is unclear how well purely sequence-based methods that rely on binding site co-occurrences can accurately detect that subset of locations which are cooperatively bound by both TFs. However, differentiating between a location that is cooperatively bound by a pair of TFs from a second location that is not cooperatively bound is possible through ChIP-seq (chromatin immuno-precipitation and sequencing) profiles of both TFs.

ChIP-seq provides a list of locations bound by a TF across the genome *in vivo*, which are referred to as *peaks*, along with peak *intensities* whose values are proportional to the TF’s affinity for the sequence bound at these locations [24]. Three sets of ChIP-seq would need to be performed to determine locations where a pair of TFs, A and B, are cooperatively bound. First, two ChIP-seq experiments are performed to determine binding locations of A and B in cells. A third ChIP-seq is performed to find binding locations of A after B is genetically knocked out. We define a location to be *cooperatively bound* by A and B if A no longer binds DNA, or has a lower peak intensity, after B is knocked out. We consider locations where A continues to bind DNA with no change in its intensity after B is knocked out to be *non-cooperatively bound*. We refer to this set of three experiments necessary to find locations where A is cooperatively bound by B as A-B, and refer to A as the *target* TF and B as the *partner* TF. Instead of knocking out B, if a ChIP-seq is performed to find binding locations of B after A is knocked out, we can infer locations where B is cooperatively bound by A. This dataset is labeled B-A, with B and A referred to as target and partner TFs, respectively. We note that this definition of cooperative binding between the target and partner TF is an operational one based on knockout data and is independent of the mechanism that generates the cooperative binding effect, of which there are several [22, 25].

However, ChIP-seq profiles of the target TF after the partner TF has been knocked out may not be easily available. To find regions where the target TF is cooperatively bound by a partner TF in the absence of such data, we propose the ChIP-seq Peak Intensity - Expectation Maximisation (CPI-EM) algorithm. At each location where ChIP-seq peaks of two TFs overlap each other, CPI-EM computes a probability that the location is cooperatively bound by both TFs. The highlight of this algorithm is that it utilizes only peak intensities to detect cooperative binding, and does not rely on binding site searches within ChIP-seq peak regions. CPI-EM relies on the observation that a target TF tends to be more weakly bound when it cooperatively bound DNA with a partner TF, in comparison to regions where it did not cooperatively bind DNA. We observed this to be the case in ChIP-seq datasets we analyzed from *E. coli*, *S. cerevisiae*, and *M. musculus* genomes [26, 1]. We chose these datasets because they included ChIP-seq data from the target TF after the partner TF had been knocked out, which allowed us to validate and measure the accuracy of CPI-EM in detecting regions where the target TF is cooperatively bound to DNA.

We compare the performance of CPI-EM with that of two other algorithms — a sequence-independent algorithm that detects cooperative binding based on the distance between the summits of ChIP-seq peaks of both TFs, and a published sequence-based algorithm, STAP (Sequence To Affinity Program) [17], that detects cooperative binding based on the binding site composition of a location. We find that CPI-EM outperforms both these algorithms. Importantly, since CPI-EM detects far more cooperative interactions amongst lower intensity ChIP-seq peaks than STAP, our work demonstrates the potential of sequence-independent algorithms such as CPI-EM to complement existing sequence-dependent algorithms in detecting more cooperatively bound locations.

## 2 Results

### 2.1 Peaks of target TFs have lower intensities when they are cooperatively bound when compared to non-cooperatively bound peaks

We inferred cooperative and non-cooperative binding using knockout data from ChIP-seq datasets of FIS-CRP and CRP-FIS pairs in *E. coli* in early-exponential and mid-exponential growth phases (accession number GSE92255), GCN4-RTG3 and RTG3-GCN4 in *S. cerevisiae* [1], FOXA1-HNF4A, FOXA1-CEBPA, and HNF4A-CEBPA in the mouse (*M. musculus*) liver [26]. A summary of the data is shown in Supplementary Table S1.

Figure 1A summarizes trends in cooperative and non-cooperative TF-DNA binding seen in these datasets. Cooperatively and non-cooperatively bound locations were determined using ChIP-seq data from genetic knockouts as discussed in Methods, with the intensity of a peak call being chosen as the 6th column of the narrowPeak output of the peak call files. We also analyzed only those ChIP-seq peaks whose peak intensities were high enough for their irreproducible discovery rate (IDR) or their false discovery rate (FDR) to be less than a specified threshold (see Supplementary Section S1). Cooperatively bound target TF peak intensities were significantly lower than those of non-cooperatively bound target TF peaks across each of the TF-TF pairs (Wilcoxon rank-sum test, *p* ≪ 0.001). In contrast, there was no consistent trend in the intensities of the partner TF in each of these pairs. We checked if these results arose from the variation in the length of the peak regions between different TFs. To control for this, we first trimmed the ChIP-seq peaks of all datasets in Figure 1A to 50 base pairs on either side of the peak summits, and then calculated anew the set of cooperatively and non-cooperatively bound regions using knockout data. We found no change in the trends seen in Figure 1A, with peak intensities of cooperatively bound primary TFs continuing to be lower than those of non-cooperatively bound primary TFs (Figure S4).

**Figure 1:**
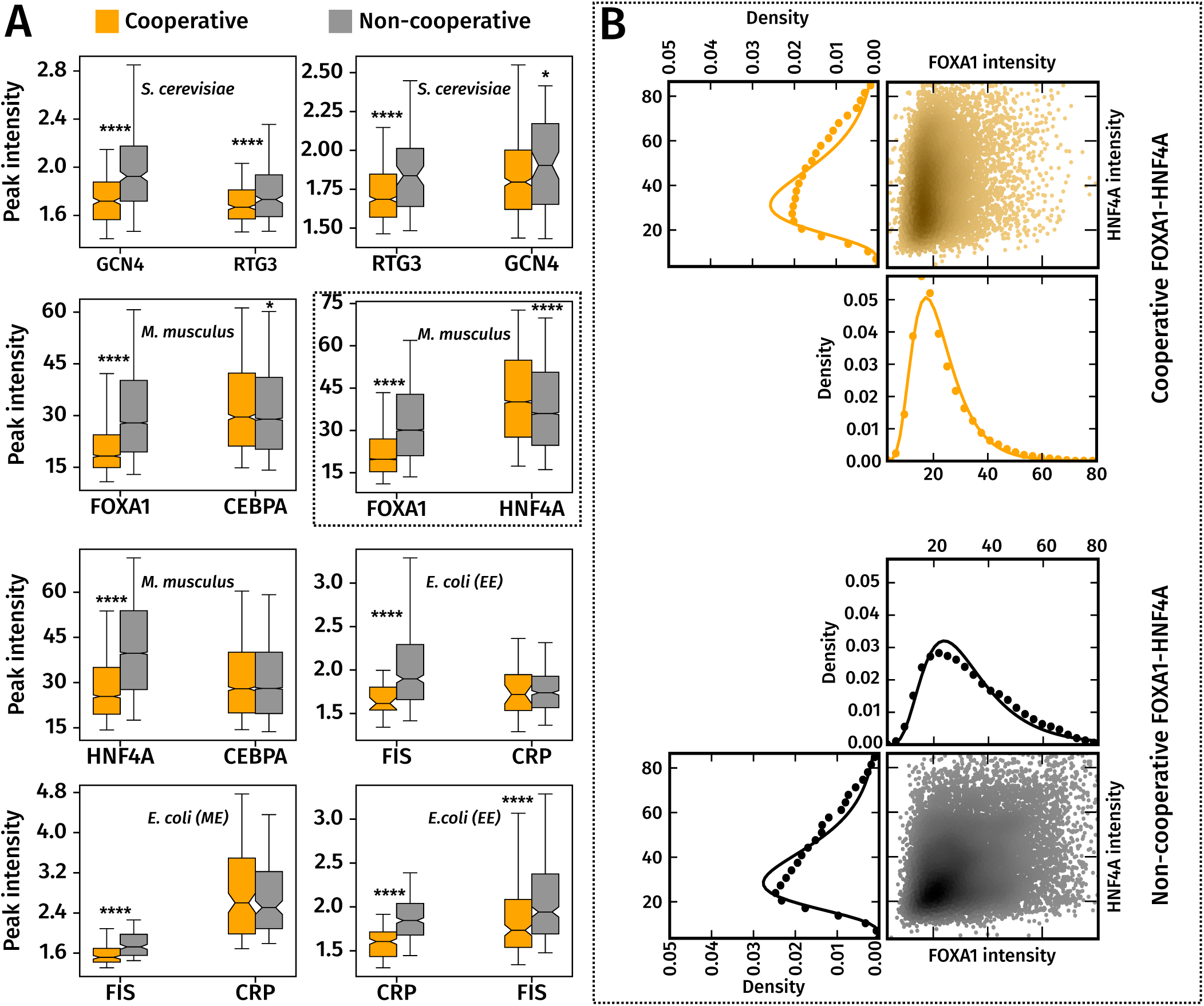
Cooperatively bound target TFs are significantly more weakly bound than non-cooperatively bound target TFs. **(A)** Box-plots of peak intensity distributions of cooperatively (orange) and non-cooperatively (gray) bound TF pairs, with target TFs on the left and partner TFs on the right. ****, *** and ** indicate p-values of < 10^−4^, 10^−3^ and 10^−2^ from a Wilcoxon rank sum test. The whiskers of the box plot are the 5th and 95th percentiles of the distributions shown. **(B) ChIP-seq peak intensity distributions can be approximated by a Log-normal distribution**. Marginal peak intensity distributions of FOXA1 and HNF4A peaks (in filled black and orange circles), with fitted Log-normal distributions (solid black and orange lines). These, and similar distributions for the other TF pairs were better approximated by a Log-normal distribution, which was evident from the higher log-likelihood value associated with a Log-normal fit, compared to a Gaussian or Gamma distribution (Supplementary Table S4). Along side the marginal intensity distributions of FOXA1 and HNF4A is a scatter plot of (FOXA1,HNF4A) peak intensity pairs from cooperatively and non-cooperatively bound regions. The scatter points are colored according to the density of points in that region, with darker shades indicating a higher density. cooperative and non-cooperative FOXA1 and HNF4A peaks are shown. The density of points in the scatter were computed using the Gaussian kernel density estimation procedure in the Python Scipy library.

We proceeded to compare the motif scores of target and partner TFs between cooperatively and non-cooperatively bound regions. We used motifs from the HOCOMOCO v10 [27] and ScerTF databases [28] for *M. musculus* and *S. cerevisiae* TFs, while we used the MEME suite [29] to determine motifs for FIS and CRP in the *E. coli* data (see Supplementary Figure S7). We calculated motif scores from the sequences underlying each ChIP-seq peak using the SPRY-SARUS scanner [27] (see section 4.5.1 in Methods). In peaks which contained multiple matches to the TF’s motif, we retained only the match that had the highest motif score for further analyses.

Similar to the trends in peak intensities in Figure 1A, we found that the motif scores of the target TF were significantly lower in cooperatively bound regions than in non-cooperatively bound regions (Supplementary Figure S3) while there was no such trend in the motif scores of the partner TF between both sets of regions. We then computed the Pearson correlation coefficient (*R*^2^) between the motif scores and intensities of peaks within each dataset and found different trends across datasets (Supplementary Figure S5). The motif scores were significantly correlated with peak intensities in the *M. musculus* datasets, but this was not the case with the remaining datasets. This means that even though the motif scores of the target TF were lower in cooperatively bound regions, they did not explain the lower target TF peak intensities observed in these regions.

Some of the peaks in these datasets may have resulted from indirect or tethered binding, where the TF being investigated does not directly bind DNA but is bound to a second protein that in turn binds DNA [30, 31, 32, 33]. If a target TF were to bind DNA indirectly via the partner TF, knocking out the partner TF would lead to a loss of the target TF’s ChIP-seq peak, or a reduction in its intensity. Such a target TF peak, which we consider cooperatively bound based on information from the ChIP-seq after the partner TF is knocked out, may, in fact, be indirectly bound.

We checked if the presence of indirectly bound peaks accounted for the trends observed in Figure 1A by removing ChIP-seq peaks of target and partner TFs that did not contain a binding site sequence for their respective TFs (see Section 4.5.2 for a full description of our method to remove indirectly bound peaks). To remove indirectly bound peaks in a single ChIP-seq experiment, we first computed the motif scores of the strongest binding site within each peak. We then computed a control distribution from motif scores of the strongest binding site within sequences that were unbound in the ChIP-seq experiment (Supplementary Figure S8). We used the 90th percentile of this control distribution as a threshold to detect indirectly bound peaks, where ChIP-seq peaks whose motif scores were lower than the 90th percentile of this distribution were declared as indirectly bound.

The removal of these peaks significantly lowered the number of doubly bound regions available for further analysis of the early-exponential phase CRP-FIS and RTG3-GCN4 datasets (see Supplementary Table S2). Nonetheless, we found that even after indirectly bound ChIP-seq peaks were removed from our analysis, cooperatively bound target TF peaks tended to have lower intensities (Supplementary Figure S2). We also found that the motif scores of the target TF in cooperatively bound peak continued to be lower than those of non-cooperatively bound target TF peaks. (Supplementary Figure S3B). The removal of the indirect peaks in the *M. musculus* dataset significantly weakens the correlation between motif scores and peak intensities, which was higher when indirectly bound peak were present in the data (Supplementary Figure S6).

Since the target TF intensity distributions from cooperatively bound regions significantly differed from those of non-cooperatively bound regions, it should be possible to accurately label a pair of overlapping peaks as cooperative or non-cooperative, based solely on their peak intensities and without carrying out an additional knockout experiment. For instance, in the FOXA1-HNF4A dataset, a FOXA1 peak that has an intensity value of 5 is ≈3.4 times more likely to be cooperatively bound with HNF4A than to be non-cooperatively bound with it. In clear-cut cases such as these, knowledge of the underlying sequence that is bound is not necessary to detect a cooperative interaction.

### 2.2 CPI-EM applied to ChIP-seq datasets from M. musculus, S. cerevisiae and E. coli

The ChIP-seq Peak Intensity - Expectation Maximisation (CPI-EM) algorithm works as illustrated in Figure 2 (with a detailed explanation in the Methods). We present a brief explanation below with each step illustrated in Figure 2B.

**Figure 2:**
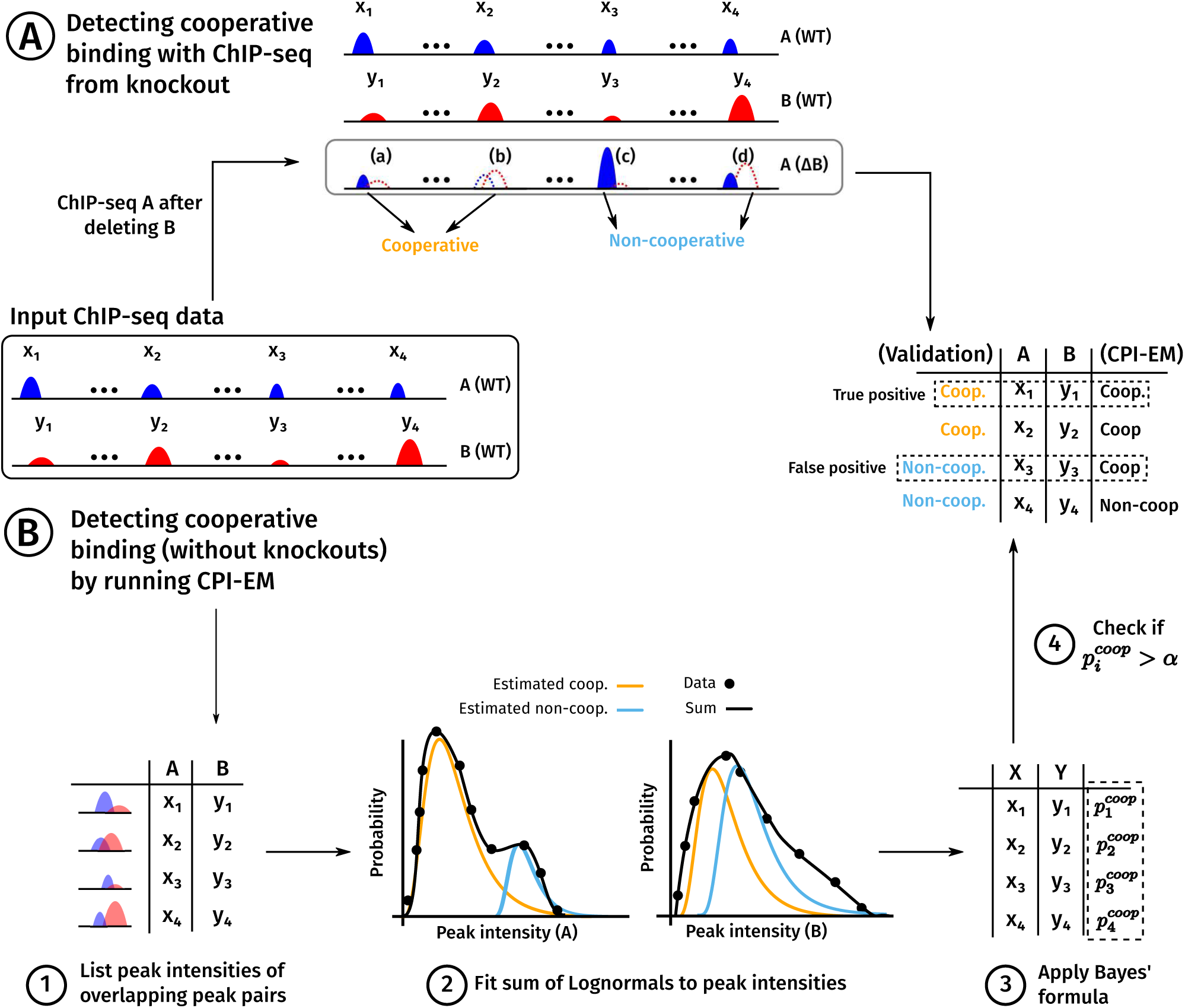
A schematic of the use of the CPI-EM algorithm and ChIP-seq from knockout data to separately identify cooperative bound transcription factor pairs. ChIP-seq experiments carried out on two TFs, A and B, yield a list of locations that are bound by both TFs, along with peak intensities at each location. From this data, there are two ways in which we find genomic locations that are cooperatively bound by A and B. (A) A method for inferring these locations from a ChIP-seq of A carried out after B is genetically deleted. Locations where a peak of A either disappears altogether, or is reduced in intensity after knocking out B are labelled as cooperatively bound. In contrast, locations where a peak of A either remains unchanged or increases in intensity are labelled as non-cooperatively bound (see section “Using ChIP-seq data from a genetic knockout to infer cooperative binding” in Methods). (B) Steps in predicting cooperatively bound locations are shown, where the numbers correspond to those in the section “The ChIP-seq Peak Intensity - Expectation Maximisation (CPI-EM) algorithm” in Methods. (1) The input to CPI-EM consists of a list of genomic locations where a peak of A overlaps a peak of B by at least a single base pair. Note that the ChIP-seq of A after B is knocked out is not an input to the algorithm. (2) Each of these overlapping intensity pairs is fit to a model that consists of a sum of two probability functions. These functions specify the probabilities of observing a particular peak intensity pair given that it comes from a cooperatively or non-cooperatively bound region. These probabilities are computed by fitting the model to the input data using the expectation-maximization algorithm (see Supplementary Section S6). (3) Bayes’ formula is applied to the probabilities computed in step (2) to find the probability of each peak intensity pair being cooperatively bound. (4) Each cooperative binding probability computed in step (3) that is greater than a threshold *α* is declared as cooperatively bound. We compare this list of predicted locations with the list of cooperatively bound locations inferred from knockout data in order to compute the number of correct and incorrect inferences made by CPI-EM.

The first step is to prepare the input to CPI-EM, which consists of a list of genomic locations where a peak of A overlaps a peak of B by at least a single base pair. Note that the genomic locations of peaks of A after B has been knocked out is not an input, since the goal of CPI-EM is to detect regions where A is cooperatively bound by B without using information from the knockout of B. In the second step, each of these overlapping intensity pairs is fit to a model that consists of a sum of two probability functions. These functions specify the probabilities of observing a particular peak intensity pair given that it comes from a cooperatively or non-cooperatively bound region. These probabilities are computed by fitting the model to the input data using the expectation-maximization algorithm (see Supplementary Section S6). In the third step, Bayes’ formula is applied to the probabilities computed in the previous step to find the probability of each peak intensity pair being cooperatively bound. Finally, each cooperative binding probability computed in the third step that is greater than a threshold *α* is declared as cooperatively bound. To validate these predictions, we compare this list of predicted locations with the list of cooperatively bound locations inferred from knockout data (Figure 2A) in order to compute the number of correct and incorrect inferences made by CPI-EM.

Figure 3 shows the result of the CPI-EM algorithm when used to predict genomic regions that are cooperatively bound by FOXA1-HNF4A, RTG3-GCN4 and FIS-CRP in *M. musculus, S. cerevisiae* and early-exponential phase cultures of *E. coli*, respectively. The top row shows histograms of the cooperative binding probabilities 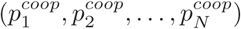, which are computed by CPI-EM, for all peak intensity pairs from each of the three datasets. The height of each bar is the fraction of peak intensity pairs in each probability bin that are actually cooperatively bound (termed true positives, which are calculated based on knockout data as explained in Methods). True positives are distributed differently between the bins across different datasets. The distribution of cooperative pairs into each of these bins determines the number of errors made when all peak pairs with *p*_*coop*_ > *α* are declared as cooperatively bound. The false positive rate (FPR) of the CPI-EM algorithm is the fraction of non-cooperatively bound regions erroneously declared as cooperatively bound, while the true positive rate (TPR) is the fraction of cooperatively bound regions that are detected. Both these quantities are functions of *α*, and are estimated as

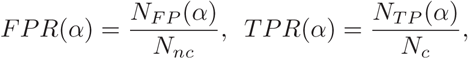

where *N*_*FP*_(*α*) is the number of non-cooperatively bound regions mistakenly declared as cooperatively bound at a threshold *α*, while *N*_*TP*_(*α*) is the number of cooperatively bound regions correctly declared as cooperatively bound with the threshold *α. N*_*c*_ and *N*_*nc*_ represent the total number of cooperatively bound and non-cooperatively bound regions, respectively, which are computed separately from the knockout data. The receiver operating characteristic (ROC) curves at the bottom row of Figure 3 shows the trade-off between *false positive rates* and *true positive rates* of CPI-EM at different values of α. A larger value of *α* results in fewer false positives in the final prediction set but also results in fewer true positives being detected. For instance, in the FOXA1-HNF4A dataset, *α* = 0.73 allows nearly 50% of all cooperative interactions to be detected. If *α* is lowered to 0.17, more than 90% of cooperative peak pairs can be detected, but there will be more false positives in this prediction set since the FPR at this value of *α* is three times higher than that at *α* = 0.73. The area under the ROC (auROC) curve provides a way of quantifying the detection performance of an algorithm. The auROC is a measure of the average true positive rate of the CPI-EM algorithm, with a higher value representing better detection performance. Thus, the auROC provides a way of comparing between different detection algorithms.

**Figure 3:**
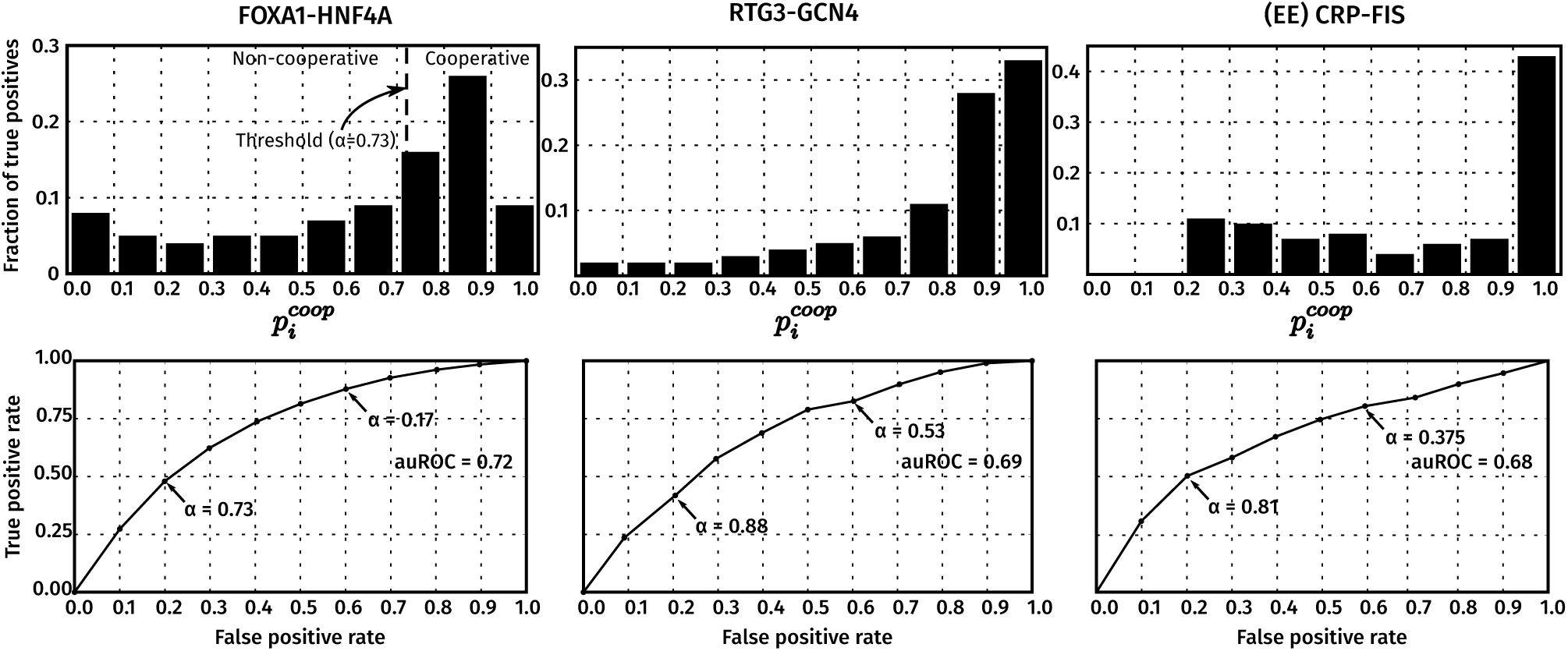
CPI-EM applied to ChIP-seq datasets from *M. musculus* (FOXA1-HNF4A), *S. cerevisiae* (RTG3-GCN4) and early-exponential phase cultures of *E. coli* (CRP-FIS). For each dataset, CPI-EM computes a list of cooperative binding probabilities at all the locations bound by the TF pair under consideration. **Top row: The fraction of cooperatively bound pairs, as determined from knockout data, that fall into each cooperative binding probability bin**. The bins are equally spaced with a width of 0.1 and the heights of the bars within each histogram add up to 1. **Bottom row: Receiver operating characteristic (ROC) curves that evaluate the performance of CPI-EM in detecting cooperatively bound pairs**. The curve is generated by calculating, for each value of *α* between 0 and 1, the true and false positive rate of the algorithm. The true positive rate (*TPR(α)*) is the ratio of the number of cooperatively bound regions detected (when *p_coop_* is compared to a threshold of *α*) to the total number of regions that are found to be cooperatively bound from the knockout data. The false positive rate (*FPR(α)*) is the ratio of the number of non-cooperatively bound regions mistakenly detected as cooperatively bound (when *p*_*coop*_ is compared to a threshold of *α*), to the total number of regions that are found to be non-cooperatively bound from the knockout data. Small values of *α* give a higher TPR, but at the cost of a higher FPR. The area under the ROC (auROC) is a measure of detection performance, whose value cannot exceed 1, which corresponds to a perfect detector. Given the auROC of two different algorithms, the one with a higher is better, on average, at detecting cooperative binding.

In the ROC curves shown in Figure 3, CPI-EM fits a Log-normal distribution to the peak intensities of the TFs in each dataset. We chose the Log-normal distribution because it gave a higher log-likelihood fit to peak intensities compared to Gaussian and Gamma distributions in most datasets (see Figure 1B and Supplementary Table S4). However, we still compared the auROC resulting from fitting a Log-normal distribution with the auROCs obtained from fitting Gamma and Gaussian distributions to peak intensities of TFs across all datasets shown in Figure 1. We found that CPI-EM with a Log-normal distribution gave the highest auROC compared to CPI-EM with Gamma and Gaussian distributions across most datasets (see Supplementary Figure S1 in Supplementary Section S4).

### 2.3 CPI-EM outperforms both STAP and a sequence-independent algorithm based on ChIP-seq peak distances in detecting cooperative binding events

Since CPI-EM relies solely on peak intensities and does not use any information from the sequences underlying ChIP-seq peaks to detect cooperative binding, we compared it with algorithms that use sequences for detecting cooperative binding. We compared CPI-EM with STAP, an algorithm which can detect genomic regions that are cooperatively bound by multiple TFs [17]. To detect cooperative binding between a TF pair A-B, where A and B are target and partner TFs respectively, STAP takes as input (a) motifs of A and B, (b) the peak intensities of A, and (c) the sequences underlying each peak of A. STAP then proceeds to build a statistical occupancy model of each sequence in order to predict peak intensities for each location, which can include cooperative or competitive interactions between A and B (see section 4.6 in Methods for more details on the inputs to STAP). STAP’s occupancy model is biophysically rigorous in that it takes into account the occurrence of multiple binding sites of A and B, binding site orientation and cooperativity between multiple copies of A and B while predicting peak intensities of the target TF. The final output of STAP is a set of predicted peak intensities for each peak of A that is input to it.

In order to detect cooperative binding, we ran STAP in two modes, which we refer to as the cooperative binding mode and the independent binding mode. In the cooperative binding mode, the occupancy model contains an extra parameter that takes into account a possible cooperative or competitive interaction between A and B. In the independent binding mode, on the other hand, the occupancy model assumes that there is no cooperative or competitive interaction that occurs between A and B. Suppose **I***ind* = {*I*_0_,*I*_1_,…, *I*_*N*_}, where *N* is the number of regions with overlapping peaks of A and B, is the set of peak intensities of A predicted by STAP when it is run in the independent binding mode, and 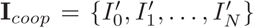 is the set of peak intensities of A predicted by STAP when it is run in the cooperative binding mode. We then define a cooperative index Δ_*j*_ for the *j - th* peak as 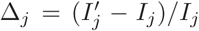, with the set of cooperative indices Δ_1_, Δ_2_,…, Δ_*N*_ constituting the region-wise predictions of cooperative binding by STAP. Locations where Δ is greater than some threshold Δ_*T*_, which could be positive or negative, are considered to be cooperatively bound.

The peak distance algorithm computes the distances between the summits of overlapping ChIP-seq peaks and declares those overlapping peak pairs whose peaks are within a threshold distance *d* to be cooperatively bound (see Section 4.4 in Methods). This detector represents a simpler sequence-independent criterion for detecting cooperative binding.

We compared the performance of STAP, the peak distance algorithm and CPI-EM (Figure 4A) in detecting cooperative interactions in the datasets shown in Figure 1, where the auROCs of CPI-EM, STAP and the peak distance detector are shown in orange, sky blue and black, respectively. We found that CPI-EM has a higher auROC than STAP in every dataset, while in the mid-exponential CRP-FIS, GCN4-RTG3 and RTG3-GCN4 datasets, STAP performed more poorly than chance. After indirectly bound peaks of target and partner TFs were removed from the input to both CPI-EM and STAP algorithms (see Section 4.5.2 in Methods), we found that CPI-EM predominantly performed better than STAP, except in the early-exponential FIS-CRP dataset where STAP had a marginally higher auROC than CPI-EM (Supplementary Figure S11A). Both STAP and CPI-EM out-perform the peak distance detector, whose auROC is lower than chance in RTG3-GCN4 and early-exponential phase FIS-CRP datasets. We encountered numerical stability issues when we ran STAP on CRP-FIS,FIS-CRP, RTG3-GCN4 and GCN4-RTG3 datasets, where the parameters of STAP’s occupancy model did not converge to the same set of parameters when we ran it multiple times (see Section 4.6.1 in Methods). These datasets are marked with an asterisk in Figure 4A.

**Figure 4:**
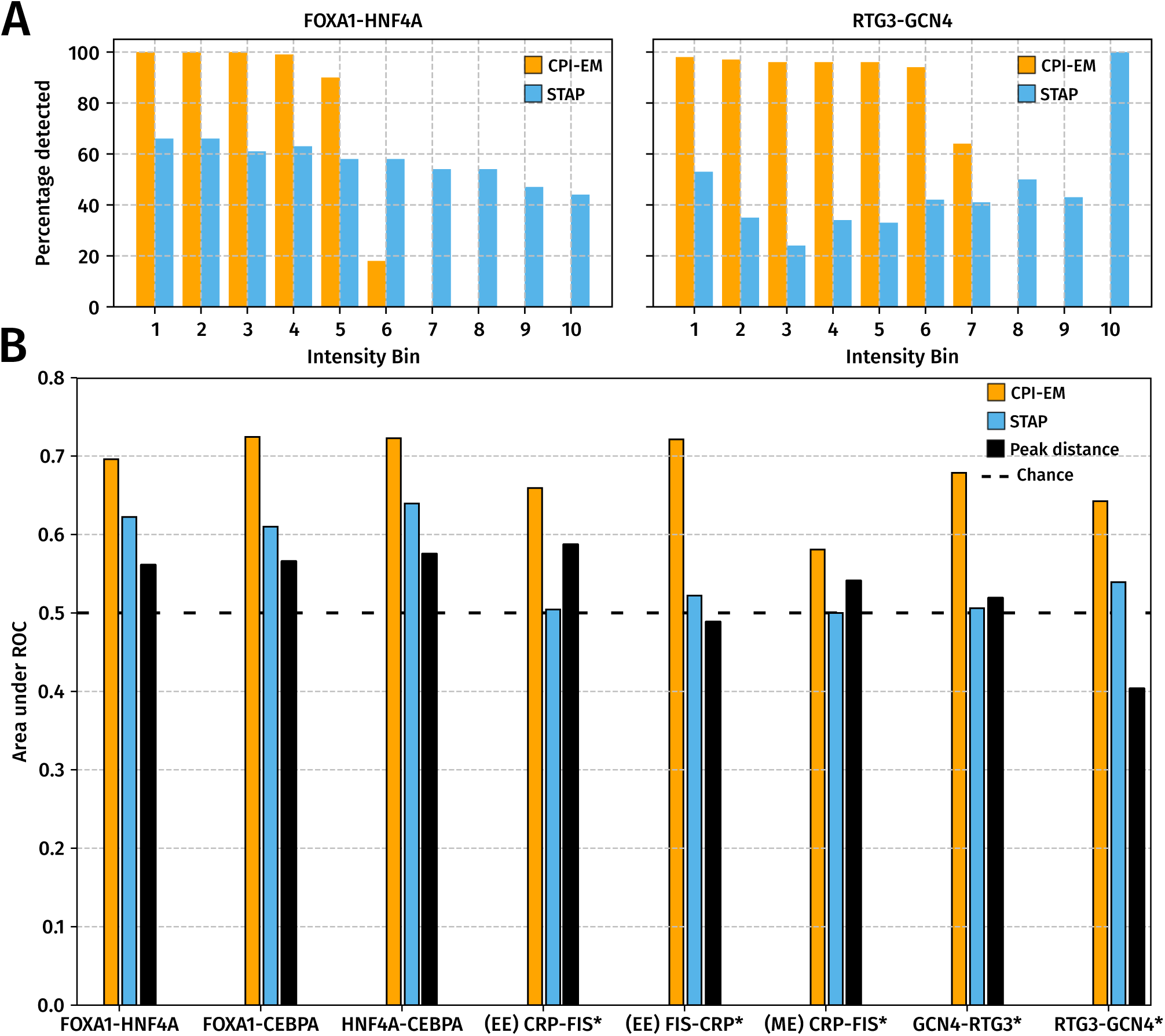
CPI-EM outperforms STAP and the peak distance detector in detecting cooperatively bound TF pairs across different datasets, even though STAP can better detect cooperatively bound target TF peaks that have high intensities. **(A)** The auROCs of CPI-EM and STAP are shown in orange and sky blue, respectively. The auROC of the chance detector, which is always 0.5 is shown by a dashed line. The datasets marked with an asterisk (*) are those where STAP was numerically unstable (see section 4.6.1 in Methods). The complete ROC curves for STAP and CPI-EM are shown in Supplementary Figure S11B and that of the peak distance detector in Supplementary Figure S9. **(B)** CPI-EM detects more cooperative interactions amongst low intensity target TF peaks but STAP detects more such interactions amongst higher intensity target TF peaks. On the *x*-axis, cooperatively bound FOXA1-HNF4A and RTG3-GCN4 peak pairs are divided into ten bins based on the intensity of the target TF, with the 10th bin having the highest intensity target TF peaks. The *y*-axis represents the percentage of cooperative peak pairs actually detected by CPI-EM (orange) or STAP (sky blue) in each bin at a false positive rate of 40%.

While CPI-EM detects more cooperative interactions than STAP at a given false positive rate, STAP detects more cooperative interactions amongst higher intensity target TF peaks than CPI-EM. This is shown in Figure 4B, we divided cooperatively bound FOXA1-HNF4A and RTG3-GCN4 peak pairs into ten bins based on the peak intensities of the target TFs in each data set, with the 10th bin containing peak pairs with the highest target TF peak intensities. In both datasets, we ran CPI-EM and STAP at thresholds that resulted in a relatively high false positive rate (~ 40%) and calculated the fraction of cooperatively bound peak pairs detected by both algorithms from each intensity bin. While CPI-EM detected nearly all cooperatively bound peak pairs from the lower intensity bins, it did not detect any cooperative interactions amongst the higher intensity bins. In contrast, STAP was able to detect cooperative interactions from each of the intensity bins, although the fraction detected within each bin was smaller compared to CPI-EM.

## 3 Discussion

Cooperative binding is known to play a role in transcription factor binding site evolution and enhancer detection [34]. Cooperativity is also known to influence cis-regulatory variation between individuals of a species [35], which could potentially capture disease-causing mutations that are known to occur in regulatory regions of the genome [36]. CPI-EM is suited to study these phenomena since it can detect instances of cooperative binding between a pair of transcription factors that may occur anywhere in the genome. While sequence-based approaches to cooperative binding detection have been proposed [10, 11, 12, 13, 17, 14, 15, 16, 1], none use ChIP-seq peak intensities as the sole criterion to detect cooperativity. We compare CPI-EM to a sequence-based approach, STAP [17], and a simpler sequence-independent algorithm based on the distance between target and partner TF peaks, and show that CPI-EM detects more cooperative interactions than either of them. However, STAP is better able to detect cooperative interactions amongst high-intensity ChIP-seq peaks. Given that CPI-EM and STAP detected interactions amongst different peak populations, this shows that sequence-independent methods like CPI-EM can usefully complement sequence-based detection algorithms.

### 3.1 Assumptions in the CPI-EM algorithm

The assumption that cooperatively bound target TFs are more weakly bound, on average, than non-cooperatively bound target TFs is the key assumption in the CPI-EM algorithm. This assumption was based on our comparison of cooperatively and non-cooperatively bound target TFs in *E. coli, S. cerevisiae* and *M. musculus* genomes. We checked if cooperatively bound TFs continue to be more weakly bound than non-cooperatively bound TFs even after indirectly bound peaks are removed from our analysis. We detected indirectly bound peaks based on a sequence-based motif analysis of the ChIP-seq peaks (see Methods) and note that there is currently no sequence-independent method to detect indirect binding in ChIP-seq data. A method like CPI-EM will declare an indirectly bound peak as cooperatively bound. However, we have shown that sequence-based criteria, such as the one employed in our analysis, or other published methods [32, 30, 31] can be used to filter out such ChIP-seq peaks before they are input to the CPI-EM algorithm. Furthermore, we show that filtering out these peaks before they are input to CPI-EM does not impact the ability of CPI-EM to detect cooperatively bound regions that are not indirectly bound (Supplementary Figure S10). However, this approach to filtering out indirectly bound peaks may discard genuine low-affinity binding sites that are actually occupied in the ChIP-seq experiment. This is because in most methods meant to detect indirect binding, a peak with a low motif score has a much higher probability of being declared as indirectly bound than a peak with a high motif score.

A caveat about the predictions of CPI-EM is that when it declares a region to be cooperatively bound by a pair of TFs, it does not implicate any particular mechanism of cooperative binding. Since CPI-EM analyzes the peak intensities of only the two TFs in question, it is in principle possible that a third TF or a nucleosome mediates the cooperative binding that is detected by CPI-EM. Thus, CPI-EM can be used to only select locations of interest that are cooperatively bound in this manner, but further computational or experimental analysis would be required to find the mechanism that give rise to the observed cooperative binding effect at each location.

Our choice of TFs to validate CPI-EM was motivated by the availability of ChIP-seq from the knockout of partner TFs in each of these datasets. The importance of data from TF knockouts arises from recent studies on cooperative binding [21, 23, 19, 20], which suggest that a pair of TFs that bind one genomic location cooperatively may not do so in a second location if there are differences in the length or the composition of the sequence linking both TF binding sites. In the absence of data from a ChIP-seq of one of the TFs after the other has been knocked out, it is impossible to ascertain which of these locations are cooperatively bound.

Our observation that a TF that cooperatively bound DNA with the help of a partner TF was more weakly bound than when it non-cooperatively bound DNA (Figure 1) is likely a signature of a short-range pair-wise cooperative interaction. For instance, the interactions between GCN4 and RTG3 were independently verified in the publication that reported this ChIP-seq data [1]. Along with the peak intensities of the target TF, the motif scores of the target TF are also significantly lower in cooperatively bound regions. However, the correlation between motif scores and peak intensities in cooperatively bound regions were low, which means that the motif scores do not directly explain the low target TF peak intensities in cooperatively bound regions. However, earlier ChIP-seq studies [37, 38] have also found a low correlation between motif score and peak intensity. These studies suggest that the correlation is increased once other factors such as chromatin accessibility have been taken into account.

Low affinity binding sites are known to be evolutionarily conserved and functionally important in the *Saccharomyces cerevisiae* genome [39], with most of these binding sites being under purifying selection to maintain their binding affinity [40]. Cooperative binding amongst such low-affinity binding sites are known to play a crucial role in animal development. The binding of Ultrabithorax (Ubx) and Extradenticle at the *shavenbaby* enhancer in *Drosophila melanogaster* embryos [41] occurs in closely spaced low-affinity binding sites to help coordinate tissue patterning. Mutations that increased Ubx binding affinity led to the expression of proteins outside their naturally occurring tissue boundaries [41]. Similarly, low-affinity binding sites that cooperatively bind Cubitus interruptus at the *dpp* enhancer (which plays a crucial role in wing patterning in *Drosophila melanogaster*) are evolutionarily conserved across twelve *Drosophila* species [42]. Cooperative binding among low-affinity transcription factor binding sites in the segmentation network of *Drosophila melanogaster* contributes to the robustness of segment gene expression to mutations [43].

### 3.2 Challenges to cooperativity detection using ChIP-seq peak intensities

There are two main computational challenges to detecting cooperative interactions using only ChIP-seq peak intensities. As stated earlier, indirectly bound ChIP-seq peaks will be declared as cooperatively bound by CPI-EM unless these peaks are checked by a sequence-dependent analysis. The second issue with CPI-EM is that as a consequence of our assumption that cooperatively bound peaks are more weakly bound than non-cooperative peaks, CPI-EM is unlikely to detect regions where the target TF is cooperatively bound to DNA, but with a high peak intensity. We found that STAP was able to detect cooperatively bound peak pairs even if the target TF was strongly bound (Figure 4B), although it detected fewer interactions in total than CPI-EM. A method that better combines the biophysically rigorous TF-DNA occupancy model of STAP with CPI-EM’s use of peak intensities might be able to detect cooperative interactions irrespective of the intensity of the target TF.

Doing away with the assumption of cooperatively bound peaks being necessarily weaker than non-cooperatively bound peaks would allow CPI-EM to detect cooperative interactions even amongst strongly bound peaks. We hypothesize that one way to accomplish this would be to take into account the high value of mutual information (MI) is expected between the binding affinities of a pair of cooperatively bound TFs [44]. The MI would then be a tenth parameter the joint probability model fit to peak intensity data (in step 2 of the CPI-EM algorithm). The precise form of such a modified joint probability model is not obvious, but it would increase the probability that a high MI peak intensity pair would be labeled as cooperative, even if the target TF were strongly bound. However, we found that the MI between the ChIP-seq peak intensities (and motif scores) of cooperatively bound TFs was low even after indirectly bound peaks were removed (Supplementary Table S3). It is possible that peak intensities obtained from experimental protocols such as ChIP-nexus [45, 46] and ChIP-exo [47, 33] might capture the high MI expected between cooperatively and non-cooperatively bound TFs. If this is indeed the case, our suggested modifications to CPI-EM would allow it detect more cooperative interactions between a pair of TFs.

The peak distance detector (Supplementary Figure S1) did not consistently detect cooperative binding across the datasets we tested it on. This detector is based on the premise that ChIP-seq peak summits that are closer together are more likely to interact with each other. The peak distance detector represented a potentially simpler criterion to detect cooperative binding compared to peak intensities. Even though TFs that were bound closer to each other were found to be more likely to interact with each other in *in vitro* studies [21, 19], the inconsistent performance of the peak distance detector shows that peak intensities are a better sequence-independent criterion to detect cooperative binding.

Ultimately, our method aims to detect cooperatively bound locations without making any direct assumptions about the genomic sequence of that location. Therefore, it provides a useful way of finding binding sequence patterns that allow for cooperative binding to occur *in vivo* but lie outside the range of existing sequence-based algorithms.

## 4 Methods

### 4.1 ChIP-seq processing pipeline

A single ChIP-seq “peak call” consists of the genomic coordinates of the location being bound, along with a *peak intensity*. We determined ChIP-seq peak locations of different transcription factors from multiple genomes, namely, *E. coli* (GSE92255), *S. cerevisiae* [1], cells from target *M. musculus* liver tissue [26]. We used our own ChIP-seq pipeline to process raw sequence reads and call peaks from *M. musculus* and *S. cerevisiae* data, and utilized pre-computed peak calls with the remaining datasets. This ensured that our validation sets were not biased by procedures employed in our pipeline. See Supplementary Section S1 for details of our ChIP-seq pipeline for processing these datasets.

### 4.2 Using ChIP-seq data from a genetic knockout to infer cooperative binding

From ChIP-seq profiles of a pair of TFs, A and B, we classified genomic regions containing overlapping ChIP-seq peaks of A and B as cooperative or non-cooperative, based on the change in peak rank of A in response to a genetic deletion of B. The ranks are assigned such that the peak with rank 1 has the highest peak intensity. In our analysis, we consider a genomic region to be doubly bound by A and B if their peak regions overlap by at least a single base pair. We used pybedtools v0.6.9 [48] to find these overlapping peak regions.

At each doubly bound genomic location, we define A as being cooperatively bound by B if (a) the peak rank of A in the presence of B is significantly higher (i.e., closer to rank 1) than the peak rank of A measured after the deletion of B, or (b) if A’s peak is absent after the deletion of B.

On the other hand, if the peak rank of A in the presence of B is significantly lower (i.e., further from rank 1) than the peak rank of A after the deletion of B, or if it stays the same, we classify this as competitive or independent binding, respectively. We refer to both these classes as non-cooperative binding. See Supplement Section S5 for details on the statistical tests we performed to detect significant changes in peak ranks of A upon the knockout of B. These tests require ChIP-seq data from multiple replicates. In the CRP-FIS, and FIS-CRP datasets, peak calls from individual replicates were not available, therefore we used only peak losses to find cooperatively bound locations in these datasets.

### 4.3 The ChIP-seq Peak Intensity - Expectation Maximisation (CPI-EM) algorithm

We describe the working of the CPI-EM algorithm in step-wise fashion below, where each of the steps is numbered according to Figure 2. In Figure 2 and in the description below, we assume that cooperative binding between TFs A and B is being studied, where A is the target TF and B is the partner TF.

**Step 1**: From the ChIP-seq of A and B, find all pairs of peaks where A and B overlap by at least one base pair. With these overlapping pairs, make a list of peak intensities (*x*_1_,*y*_1_), (*x*_2_, *y*_2_)…(*x*_*n*_,*y*_*n*_), where *x*_*i*_ and *y*_*i*_ are the peak intensities of the *i - th* peak of A and B, respectively. This list of peak intensity pairs is the input data for the CPI-EM algorithm.

**Step 2**: To this input data, fit a model of the joint probability *p(x, y*) of observing the peak intensity *x* and *y* from TFs A and B, respectively, at a given location. Our model consists of a sum of two probability functions, which are the probability of observing intensities *x* and *y* if they were (a) cooperatively bound, or (b) non-cooperatively bound. We assume that both probability functions that are fitted have a Log-normal shape. This shape is characterized by four parameters — a mean and a variance of the A and the B axes (we also examine other shapes such as the Gamma or Gaussian functions — see Supplementary Table S4). A final ninth parameter sets the relative weight of the two probability functions, which determines the fraction of overlapping pairs that are cooperatively bound. We find the best fit for these nine parameters using a procedure called expectation maximization (described in detail in Supplementary Section S6).

We make two other assumptions in this step, each of which is discussed further in Supplementary Section S6.

- The peak intensities of A and B at a location are statistically independent, irrespective of whether A and B are cooperatively or non-cooperatively bound. We found this to be a reasonable assumption after we measured the mutual information between peak intensities of A and B from cooperatively and non-cooperatively bound locations (Supplementary Table S3). Mutual information is known to be a robust measure of statistical dependence [49].
- A target TF that is cooperatively bound to DNA is, on average, bound weaker than a non-cooperatively bound target TF. We found this assumption to hold across all the datasets on which we ran CPI-EM (see section “Peak intensities of cooperatively bound target TFs are weaker than non-cooperatively bound target TFs” in Results, and Figure 1).

**Step 3**: Given the best-fit parameters, use Bayes’ formula to calculate the probability for each overlapping pair of ChIP-seq peaks to be a site of cooperative binding (see Supplementary Section **S6**).

**Step 4**: Choose a threshold probability *α* and label an overlapping pair as cooperatively bound if the probability calculated in step 3 is greater than α, and as being non-cooperatively bound otherwise. Validate with a list of known cooperative binding sites, e.g., derived from the ChIP-seq of A after B is knocked out (as described in the previous section).

### 4.4 Peak Distance Detector

For each peak intensity pair in the input data, the peak distance detector calculates the distance between the *summits* of A and B peak regions. The summit is a location within each peak region that has the highest number of sequence reads that overlap it, and is typically the most likely site at which the TF is physically attached to DNA. The peak distance detector declares doubly bound regions as cooperatively bound if the distance between peaks of A and B is lesser than a threshold distance *d.* We ran this detection algorithm on all the datasets on which CPI-EM was employed to detect cooperative binding. Our goal in using this algorithm was to determine whether the distance between peaks is a reliable criterion to discriminate between cooperative and non-cooperative binding.

### 4.5 Sequence-based analyses of ChIP-seq data

#### 4.5.1 Motif discovery and scanning

The motifs of FOXA1, HNF4A and CEBPA in *M. musculus* ChIP-seq data were sourced from the HOCOMOCO v10 database [27]. The motifs of GCN4 and RTG3 were sourced from the ScerTF database [28]. See Figure S7 for all the motifs used in our analysis.

The motifs of CRP and FIS in the wild-type, Δcrp and Δfis backgrounds were learned de novo using the MEME suite (v4.12.0) [29]. For each of these ChIP-seq datasets, we sorted the peaks according to their peak intensity and short-listed the sequences in the top 200 peaks as inputs to the MEME suite. MEME was run on these peak sequences with the options (**-bfile <genome background file>-dna-p 7-revcomp**) to generate the CRP and FIS motifs shown in Figure S7. The genome background file was created by running the **fasta-get-markov** tool of the MEME suite with default options, which created a zeroth-order Markov model of the genome.

In order to scan ChIP-seq peaks for motif matches, we used the program SPRY-SARUS [27] (http://autosome.ru/chipmunk/) with the option besthit so that only the motif with the highest match score was output for each ChIP-seq peak.

#### 4.5.2 Detecting indirectly bound peaks in a ChIP-seq dataset

In order to detect indirectly bound peaks in each ChIP-seq dataset, we first extracted a set of *N* unbound sequences, each of length *l* from the genome, where *N* is the number of peaks in the dataset and *l* is the mean ChIP-seq peak length. In RTG3, GCN4, CRP and FIS datasets, where the number of peaks was small, we created a set of 10000 unbound sequences of length *l*. We refer to this set of unbound sequences as the *negative control* dataset.

We then used the motif of the respective TF being probed using ChIP-seq and computed the score of the best motif match in each sequence of this negative control set using SPRY-SARUS as mentioned in the previous section. The distribution of the resulting set of motif scores is shown by the dashed lines in the panels of Figure S8.

The 90th percentile of this distribution, which we denote as *T*, is shown by a vertical gray line in each panel. We consider a ChIP-seq peak to be indirectly bound if the highest motif match score within the sequence of the peak is less than *T*. The solid line in each panel of Figure S8 is the distribution of motif scores from the sequences underlying the ChIP-seq peaks. The numbers in the top-right of each panel denote the number of directly bound peaks and the total number of peaks in the dataset.

This criterion for detecting indirectly bound peaks is similar to the one employed in an earlier analysis of ENCODE data [32]. In that analysis, a peak in a ChIP-seq for TF A whose sequence does not contain a subsequence that matches the motif for A but matches that for a different TF B is considered to be indirectly bound. In our case, where we are interested in detecting peaks that indicate cooperative binding of A by B, if we find that a peak of the TF A does not have a motif match whose score is above *T*, we do not search the sequence for a motif match for B but simply discard the peak altogether. This gives us the advantage of ensuring that peaks where A may be cooperatively bound by a third TF, say C, whose ChIP-seq data is not available to us, are also removed from the dataset.

#### 4.6 Detecting cooperative binding with Sequence to Affinity Prediction (STAP)

We ran STAP v2 (https://github.com/UIUCSinhaLab/STAP) to detect cooperatively bound regions across the genome. There are three inputs required to run STAP when using it to detect cooperative binding between A (target TF) and B (partner TF) —

- A training set that consists of a mixture of bound and unbound sequences from the ChIP-seq of the target TF along with their peak intensities. We followed the same procedure to construct this training set as described inthe original STAP publication [17]. We constructed this set using sequences of the 500 highest intensity peaks that were cooperatively bound (as detected from the knockout) and also 500 sequences from unbound genomic regions. Each unbound sequence was of length equal to the average length of a ChIP-seq peak in that dataset. In cases where the number of cooperatively bound peaks were less than 500, we chose upto half of the total number of cooperatively bound peaks and used sequences from non-cooperatively bound peaks to create the set of 500 bound sequences. We set the peak intensities of the bound sequences to be the score column of the peak call file (which is typically the 5th column of the peak call file), while the peak intensities of the unbound sequences were set as 0. This was in line with the
- A test set that consisted of the remaining bound sequences from ChIP-seq peaks of the target TF A that were not present in the training data.
- A motif file for the target and partner TFs being analyzed. When we ran STAP in the independent binding mode, we passed the motif of only A as an input, and when we ran STAP in a cooperative binding mode, we passed the motifs of both A and B as inputs.

As stated in the main text, we ran STAP in cooperative and independent binding modes and defined a cooperative index Δ_*j*_ for the *j - th* peak in the test dataset as 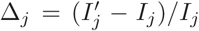, where *I*_*j*_ is the predicted peak intensity of A when there is no cooperative interaction assumed between A and B and 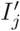 is the predicted peak intensity of A when a cooperative interaction is assumed to exist between A and B. The set of cooperative indices Δ_1_,Δ_2_, …, Δ_*N*_ constitute the region-wise predictions of cooperative binding by STAP. Locations where Δ is greater than some threshold Δ_*T*_, which could be positive or negative, are considered to be cooperatively bound. By varying Δ_*T*_, we compute the ROC of STAP (see Supplementary Section S7).

#### 4.6.1 Numerical stability of STAP runs

We found that on some datasets, particularly *S. cerevisiae* and *E. coli* datasets, STAP tended to generate different predicted peak intensities when run multiple times. To deal with such instances, we ran STAP five times each in both independent and cooperative binding modes on each dataset.

The key model parameters computed by STAP that allow it to predict peak intensities for each input sequence are the Boltzmann weights of the configuration at each sequence [17]. The Boltzmann weights computed by STAP for each sequence represent un-normalized probabilities of finding the sequence in either a bound state or an unbound state. The default diagnostic output of STAP includes the largest pair of Boltzmann weights calculated by it. Across each of the five runs of STAP, we stored this pair of Boltzmann weights and computed the coefficient of variation of each of these weights (i.e. the ratio of the standard deviation to the mean). For datasets where this coefficient of variation was greater than 10%, we considered STAP to be numerically unstable. Additionally, since Boltzmann weights represent un-normalized probabilities, they should always be non-negative. In datasets where the maximum Boltzmann weights output by STAP were negative in one of the runs, we considered STAP to be numerically unstable.

In cases where the STAP predictions differed between multiple runs, we chose that STAP run with the maximum *R*^2^ value between the predicted peak intensities and actual peak intensities as the representative one for computing the ROC curve.

## Acknowledgements

We thank Aswin Sai Narain Seshasayee, Parul Singh, Sridhar Hannenhalli, Vijay Kumar, Deepa Agashe, and Leelavati Narlikar for discussions.

## Author Contributions

V.D. conceived the study, and designed and implemented the CPI-EM algorithm. V.D., R.S., and S.K. analyzed and interpreted the results, and wrote the manuscript.

